# GenomeFace: a deep learning-based metagenome binner trained on 43,000 microbial genomes

**DOI:** 10.1101/2024.02.07.579326

**Authors:** Richard Lettich, Robert Egan, Robert Riley, Zhong Wang, Andrew Tritt, Leonid Oliker, Katherine Yelick, Aydın Buluç

## Abstract

Metagenomic binning, the process of grouping DNA sequences into taxonomic units, is critical for understanding the functions, interactions, and evolutionary dynamics of microbial communities. We propose a deep learning approach to binning using two neural networks, one based on composition and another on environmental abundance, dynamically weighting the contribution of each based on characteristics of the input data. Trained on over 43,000 prokaryotic genomes, our network for composition-based binning is inspired by metric learning techniques used for facial recognition.

Using a task-specific, multi-GPU accelerated algorithm to cluster the embeddings produced by our network, our binner leverages marker genes observed to be universally present in nearly all taxa to grade and select optimal clusters of sequences from a hierarchy of candidates.

We evaluate our approach on four simulated datasets with known ground truth. Our linear time integration of marker genes recovers more near complete genomes than state of the art but computationally infeasible solutions using them, while being over an order of magnitude faster. Finally, we demonstrate the scalability and acuity of our approach by testing it on three of the largest metagenome assemblies ever performed. Compared to other binners, we produced 47%-183% more near complete genomes. From these datasets, we find over the genomes of over 3000 new candidate species which have never been previously cataloged, representing a potential 4% expansion of the known bacterial tree of life.

## 1 Introduction

Metagenomic assembly, the simultaneous assembly of entire microbial communities, introduces additional challenges beyond those already encountered in the assembly of individual isolate genomes. Factors such as varying abundance of species, repetitive regions, low sequencing coverage, sequencing errors, the presence of multiple strains of a single species, horizontal gene transfer, and highly conserved or homologous sequences co-located in multiple species all cause ambiguities during the assembly process.

In practice, such issues inevitably lead to metagenome assemblers producing a mixture of fragmented genomes, each dispersed across an initially unknown number of contigs rather than an ideal single scaffold per genome.

To deconvolve and group these assembled sequences by their taxa of origin, tools known as metagenome binners are used. These typically work by applying clustering algorithms with per-sample sequence abundance and/or features derived from their composition, most often via some form of ‘genomic signature’ that quantifies the relative frequency of oligonucleotide patterns within the sequences [1]. Popular metagenomic binners, such as MetaBAT 2 [2] and VAMB [3], have utilized sequences’ normalized Tetra-Nucleotide Frequencies (TNF) as a way to discriminate between genomes based on the composition of the sequences themselves.

The design and effectiveness of such techniques are rooted in the observation that oligonucleotide signatures, such as Tetra-Nucleotide Frequencies, tend to remain consistent throughout genomes. Notably, larger variations exist between genomes, correlating with phylogenetic distance. While the exact reasons for this are not fully understood, it is believed that the phenomenon is mediated by species-specific replication and repair machineries, evolutionary pressures, and to a lesser extent, environmental factors [4].

Recent studies have highlighted the advantages of large-scale metagenomic coassembly [5–7]. By aggregating multiple metagenomic samples from the same environment, coassembly provides a more holistic understanding of microbial communities compared to the separate assembly of each sample, commonly known as ‘multiassembly’. This approach not only aids in detecting community members with lower sequencing depths but also contributes to the reconstruction of more complete genomes, offering a broader and more detailed view of the environment’s taxonomic diversity. While coassembly improves microbial representation, it also significantly increases the computational complexity and precision needed in downstream binning processes, due to the larger number of sequences and the variety of genomes present.

This paper introduces GenomeFace, a binner uniquely tailored for large-scale coassemblies. It was motivated by the challenges of distinguishing between numerous genomes with high precision, integrating marker genes at scale, and efficiently processing assemblies that comprise thousands of genomes and millions of sequences.

Previous research has suggested that generalizing TNF to oligonucleotide signatures with varying lengths, degenerate alphabets, and normalization methods may provide better or complementary signals for metagenome binners, and proposed further investigation into maximizing and unifying their discriminative power [4]. To this end, we draw inspiration from the field of computer vision and facial recognition to train a deep, densely connected neural network on simulated contigs from over 43,000 prokaryotic reference genomes using oligonucleotide frequencies of various lengths as input features. The network employs a softmax cross-entropy loss, incorporating modifications from CurricularFace [8] designed to enhance the discriminative ability of facial embeddings. Although we train the network as a classification problem, its ultimate goal is to produce an embedding that minimizes the distance between sequences from the same genome and maximizes the distance between sequences originating from different genomes, as to yield a robust metric for measuring the taxonomic similarity of sequences.

This style of training is typical for state-of-the-art networks used for facial recognition, however the methodology is not intrinsically linked to the task. Although initial practitioners of such techniques did not provide theoretical justification, it has been shown that minimizing cross-entropy loss can be seen as an approximate bound-optimization algorithm for minimizing a pairwise distance loss on a hyper-sphere [9], maximizing inter-class and minimizing intra-class variance. This insight forms the foundation for our neural networks, enabling us to train on tens of thousands of genomes without encountering the issues associated with more conventional metric learning loss functions at such a large scale. These problems fundamentally arise from attempting to learn directly on the distances between embeddings of sample pairs, due to the combinatorial explosion in the number of such pairings.

To group the sequences by taxa, we introduce a novel algorithm that leverages our embeddings alongside observed universal single copy genes, which are present in nearly all microbial life, to aid in the formation and selection of sequence clusters as a form of transductive learning.

Other metagenome binners have attempted to integrate universal single copy genes. For example, DASTool, an ensemble binner, takes the output of multiple metagenome binners and uses single copy marker genes to estimate the quality of each bin. At each iteration, it greedily selects an optimal bin to output, updating the non-selected candidate bins by removing sequences that occur in the greedily selected bin and then recalculating their score. MetaBinner and Semibin 2, on the other hand, greedily extracts bins that meet quality requirements, then reclusters the remaining sequences.

In contrast to the straightforward approach of using marker genes as a post-clustering oracle, our binner integrates them in a structured and well-defined way directly into the cluster formation process. Employing the embeddings and associated similarity metric produced by our neural networks, we construct hierarchy of clusters delineating various partitionings of the input sequences, finely gradating from all sequences belonging to a single genome or cluster, to each sequence being its own cluster. From this hierarchy, we precisely extract the non-redundant set of clusters that maximizes a well defined objective function that evaluates the perceived quality of the resulting genome bins. Our clustering algorithm’s kernel is based on NVIDIA’s implementation of the HDBSCAN, modified to efficiently integrate both of our embeddings and enable multi-GPU acceleration.

We demonstrate the outperformance of state-of-the-art binners on four of the simulated CAMI-II Human Microbiome datasets. Of no less importance, we showcase our binner’s ability to scale by performing three case studies on large-scale co-assemblies of complex communities, each comprising thousands of species and assembled on the Summit supercomputer from trillions of base pairs of reads. This demonstrates its efficacy in capturing the substantial taxonomic diversity that necessitates such large-scale analyses, a feat recent state-of-the-art but more computationally expensive binners are not able to reach.

The significance of the ability to proce ss these types of data sets is immense. In our flagship case study analyzing the brackish wetlands of Twitchell Island, California, GenomeFace was able to produce 1,923 medium quality Metagenome Assembled Genomes spanning 69 phyla and candidate-phyla from a single assembly, each representing a unique species, with 90 percent of these species not previously cataloged. For perspective, we compare our analysis of only Twitchell Island’s wetlands to two of the largest consorted efforts to study aquatic communities ever performed, *Tara Oceans* and *A genome catalogue of lake bacterial diversity and its drivers at continental scale* [10], both of which encompass microbial communities sampled from a diverse range of geographic locations. These studies documented 1,888 and 1,008 prokaryotic species, representing 29 and 22 phyla, respectively.

Furthermore, from our three case studies, we discover over 3,000 new candidate species not previously cataloged in the Genome Taxonomy Database, marking a 4% expansion of the known bacterial tree of life.

The successful application of GenomeFace to these extensive co-assemblies stands as a proof of principle, signifying an advance in our ability to study the full taxonomic diversity of microbial ecosystems. This includes shedding light on microbial ‘dark matter’–low-abundance organisms that have remained largely elusive and poorly understood–thereby enabling a path forward to further understand ecological interactions, nutrient cycles, the roles of microorganisms, and microbial ecology as a whole.

## 2 Results

We present the results of GenomeFace with two widely used metagenome binning tools, VAMB and MetaBAT 2, on real, large-scale multi-terabase coassemblies. We also compare GenomeFace to SemiBin 2 on small ‘Toy’ CAMI II Human Microbiome datasets.

We start by outlining our methodology and the three real datasets we used. The real datasets are metagenomes from Twitchell Island’s wetlands, Harvard Forest’s soil, and Puerto Rico’s tropical forest soil. We examine the assembly and binning processes for each case study and discuss any noteworthy observations. We excluded SemiBin from the real dataset comparison, as it failed to run.

For the simulated datasets, we examine GenomeFace’s ability to generalize by retraining the neural network for each dataset, excluding related taxa up to the family level in our training set. We test its efficacy for novel and unseen genomes.

### 2.1 Real Datasets

#### Evaluation

In each of our three case study datasets, we compared GenomeFace with VAMB and MetaBAT 2. The contamination and completeness of each genome bin produced by these binners were evaluated using CheckM2 [11]. Taxonomic assignment was conducted using GTDBTKv2 [12], with the database version being 214.0. All binners were run with a minimum output bin size of 200 kilobases. Semibin was excluded from this comparison because it failed to run.

In Figure 2(c), we compare the number of medium quality genomes produced by each binner, as defined by the Minimum Information about a Metagenome-Assembled Genome (MIMAG) standards, which specify ≥50% estimated completeness and ≤10% estimated contamination. Figure 2(a) presents a comparison of the number of near-complete genomes generated by each binner, defined as having ≥90% estimated completeness and ≤5% estimated contamination. This criterion corresponds to the MIMAG standard for high-quality genomes but omits checks for RNA gene presence [13].

**Fig. 1:**
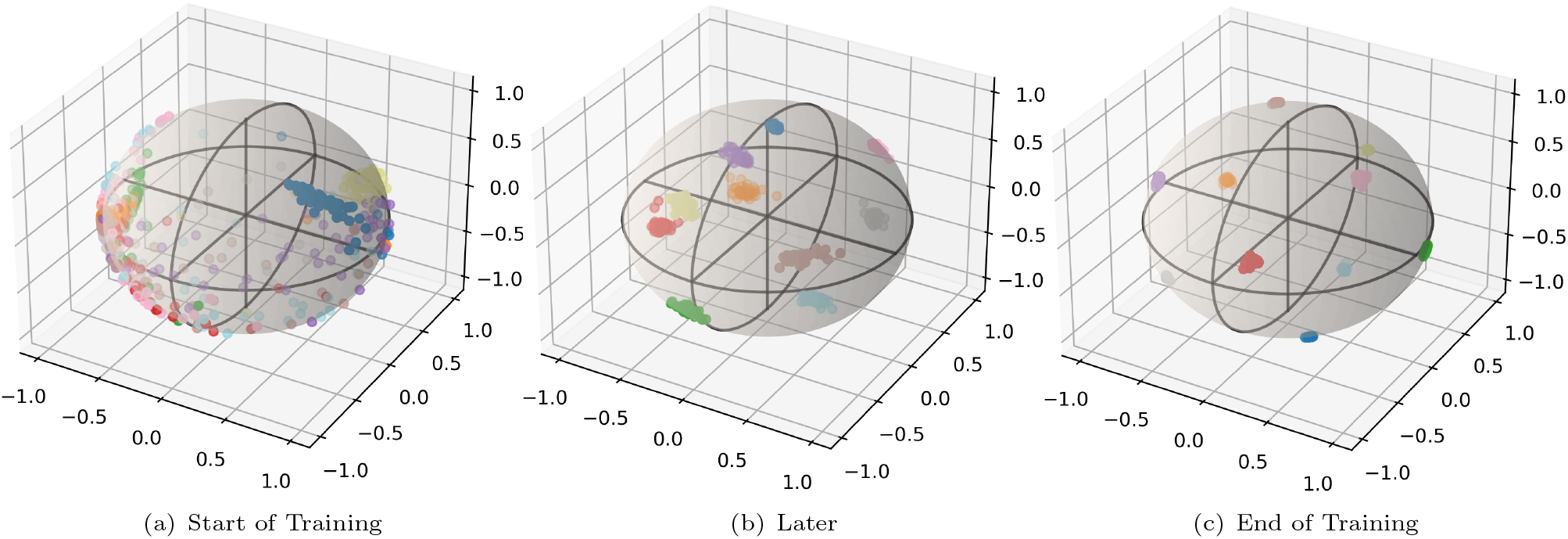
Small scale example showing the training process of our neural network, training to classify classify contigs from 10 genomes. Each point represents a contig of length 5000 from a genome being trained on, and colored based upon genome they were drawn from. The spatial position is the embedding given by our neural network, which was modified to produce embeddings in ℝ^3^ rather than ℝ^512^ for illustrative purposes. At the end of training, the sequence embeddings exhibit high intergenome and low intragenome variance

**Fig. 2:**
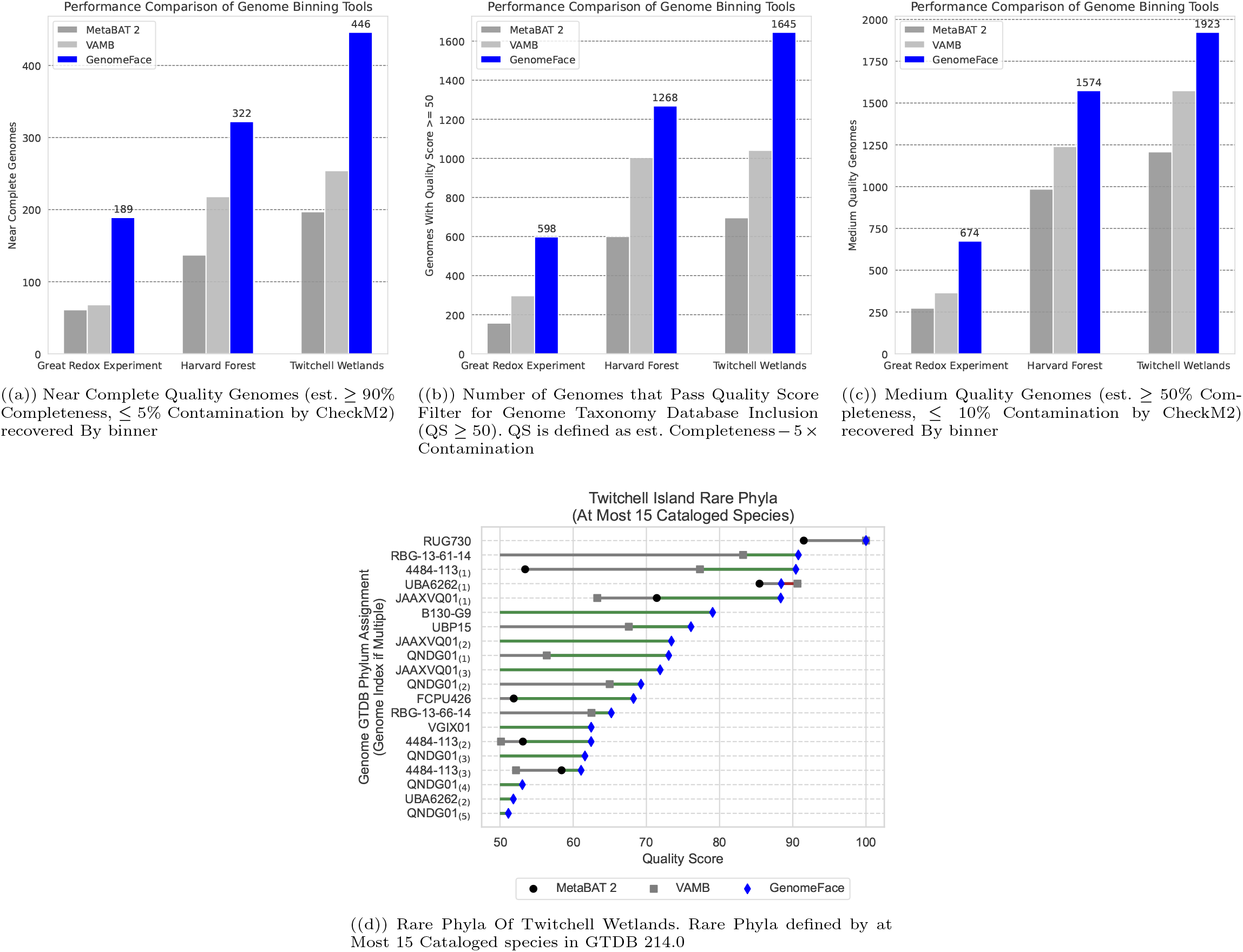
Analysis of Real Datasets

##### Case Study 1: Twitchell Island, California Wetlands

In Case Study 1, we present our findings from the 2.6 terabase coassembly of the brackish wetlands of Twitchell Island, located in the Sacramento Delta, California [14].

Utilizing MetaHipMer2 (MHM2), a distributed metagenome assembler [5], we co-assembled 2.6 terabytes of FASTQ reads from 21 samples collected across seven different locations with varying salinity and conditions [15]. This process resulted in 71.3 million scaffolds, each at least 500 bp, totaling 71.7 Gbp.

For binning with MetaBAT2, the 2.9 million scaffolds that were at least 2.5 kbp in length were used (subsequently, scaffolds of ≥ 1000 bp were attempted to be integrated post-binning), producing 3,901 bins larger than 200 kilobases, of which 1,208 were of medium quality or better. Employing VAMB with its manuscript default of sequences ≥ 2000 bp in length resulted in 1,240 medium-quality metagenome-assembled genomes (MAGs). GenomeFace was tested with its default setting for scaffolds greater than or equal to 1500 bp in length, and produced 1,923 medium quality genomes, with 68 unique phyla represented. In comparison, MetaBAT 2 generated 1,208 medium quality genomes across 66 unique phyla, while VAMB yielded 1,240 medium quality genomes, identifying 63 unique phyla. When considering near-complete genomes, GenomeFace identified 446 genomes, encompassing 46 unique phyla. MetaBAT 2 assembled 197 near-complete genomes, covering 38 unique phyla, and VAMB compiled 254 near-complete genomes, spanning 32 unique phyla.

##### Case Study 2: Soil at Harvard Forest (Barre Woods Site)

To demonstrate our method’s efficacy in characterizing microbes within a complex soil environment, we coassembled 28 metagenome samples, totaling 1.3 terabase pairs (Tbp), sequenced at the DOE Joint Genome Institute. This sequencing was part of a proposal titled “Molecular mechanisms underlying changes in the temperature-sensitive respiration response of forest soils to long-term experimental warming” [16]. The study aimed to elucidate the impact of climate change on the metabolic activities of microbial communities involved in the decomposition of soil organic matter (SOM) at the Barre Woods Harvard Forest in Petersham, MA, USA, a site of Long-Term Ecological Research (LTER) [17].

Coassembly using MetaHipMer2 (MHM2) yielded 70.8 million scaffolds, each greater than 500 bp, cumulatively amounting to 73.03 Gbp of assembled sequence.

For binning, MetaBAT 2 was employed with its default contig length settings, consistent with the approach used in Case Study 1. Similarly, VAMB was run with its manuscript default settings, as in Case Study 1. In contrast, GenomeFace was run with a minimum contig length of 2000 bp due to GPU memory requirements. GenomeFace yielded 1,574 medium quality genomes, MetaBAT 2 produced 985, and VAMB resulted in 1,204. Interestingly, all three binners identified 23 unique phyla in the medium quality genomes. For near-complete genomes, GenomeFace identified 322, encompassing 19 unique phyla. In comparison, MetaBAT 2 found 137 near-complete genomes across 17 unique phyla, and VAMB uncovered 218 near-complete genomes, identifying 15 unique phyla.

##### Case Study 3: Tropical Forest Soil

In this case study, we focus on the analysis of a substantial dataset derived from the Luquillo Experimental Forest (LEF) in Puerto Rico, a renowned site within the Long Term Ecological Research Network, as first detailed in *Riley et al* [7]. This dataset comprises an impressive 3.4 terabase pairs (Tbp) of metagenomic sequence data obtained from 95 metagenome samples.

Using 512 nodes on the Oak Ridge National Laboratory (ORNL) Summit supercomputer and taking approximately 1 hour and 24 minutes, the data were assembled into 55,342,847 scaffolds, each at least 500 bp in length.

Upon initially inspecting the results from this study, we conducted an analysis to understand the unexpectedly low number of genomes relative to the number of single-copy genes observed in the assembly. We noted that the read mappings exhibited significantly higher inter-sequence variance than expected, coupled with substantial read duplication, likely a result of PCR amplification. To mitigate this variance, we employed samtools to remove duplicate reads. However, higher variance in sequencing depth still remained. To counteract this, we enabled an auto-balancing mode in GenomeFace which mixes our two modalities using Bayesian optimization by optimizing the cluster reward. See section 4.4

Due to the larger number of metagenome samples, neither MetaBAT 2 nor VAMB were able to process on the datasets using their default settings and finish within the 12 hour walltime limit, and genomeface, and GenomeFace was constrained by GPU memory limitations. Therefore, as in the original study [7], we set all binners to use a minimum sequencing length of *≥* 3000.

In this dataset, GenomeFace yielded 674 medium quality genomes, while MetaBAT 2 produced 273, and VAMB resulted in 365. Remarkably, GenomeFace identified 29 unique phyla in the medium quality genomes, surpassing the diversity captured by MetaBAT 2 and VAMB, which identified 26 and 21 unique phyla, respectively. For near-complete genomes, GenomeFace continued to lead with 189 genomes, encompassing 19 unique phyla. MetaBAT 2, in this category, found 61 near-complete genomes spanning 15 unique phyla, and VAMB uncovered 68, identifying 14 unique phyla. Using the default mixture method, GenomeFace recovered 541 Medium Quality Genomes, and 156 near Complete genomes.

### 2.2 Simulated Datasets

In order to objectively measure the capabilities of GenomeFace, we utilized a set of simulated datasets from the Critical Assessment of Metagenomic Interpretation (CAMI) challenge [18]. CAMI’s synthetic metagenomic datasets are previously unused in benchmarking VAMB [3] and SemiBin2 [19].

We compared GenomeFace’s performance against VAMB, MetaBAT 2, and the relatively new SemiBin2. We quantified binner performance by the number of Near-Complete (NC) genomes reconstructed with a benchmark of ≥ 90% recall and ≥ 95% precision. Measurements were performed using VAMB’s command line benchmark tool, which follows the same methodology described in AMBER [20], used in the CAMI challenge [18].

A distinctive feature of GenomeFace is its use of supervised training, integrating prior knowledge into the binning process. This introduces questions about its ability to generalize across unseen taxonomic groups. To address this, we adopted a “Family Holdout” cross-validation approach.

For each of the four simulated datasets, we identified all included genomes. Subsequently, we removed any genome from the training set that was taxonomically related to any genome within the simulated datasets, based on classifications from the Genome Taxonomy Database. For example, if the airways dataset included a bacterium from the Pseudomonadaceae family, all genomes from that family would be excluded from the training set, decreasing the dataset size from around 43,000 to approximately 30,000. This exclusion strategy was executed with two distinct objectives: firstly, to verify that GenomeFace’s learning mechanism is truly understanding and applying a generalized metric for distinguishing taxa, rather than simply memorizing the training set; and secondly, to ensure that GenomeFace’s proficiency extends to categorizing taxa without any prior training on that family, showcasing its capacity to identify and classify entirely unfamiliar genomic structures.

#### Binning Performance

Across all four datasets, GenomeFace consistently demonstrated superior performance, achieving the highest number of Near Complete genomes (Figure 3(a)), performing almost identically under the Family Holdout constraint. Such results are indicative not only of GenomeFace’s precision but also of its resilience and adaptability when presented with unfamiliar taxa.

**Fig. 3:**
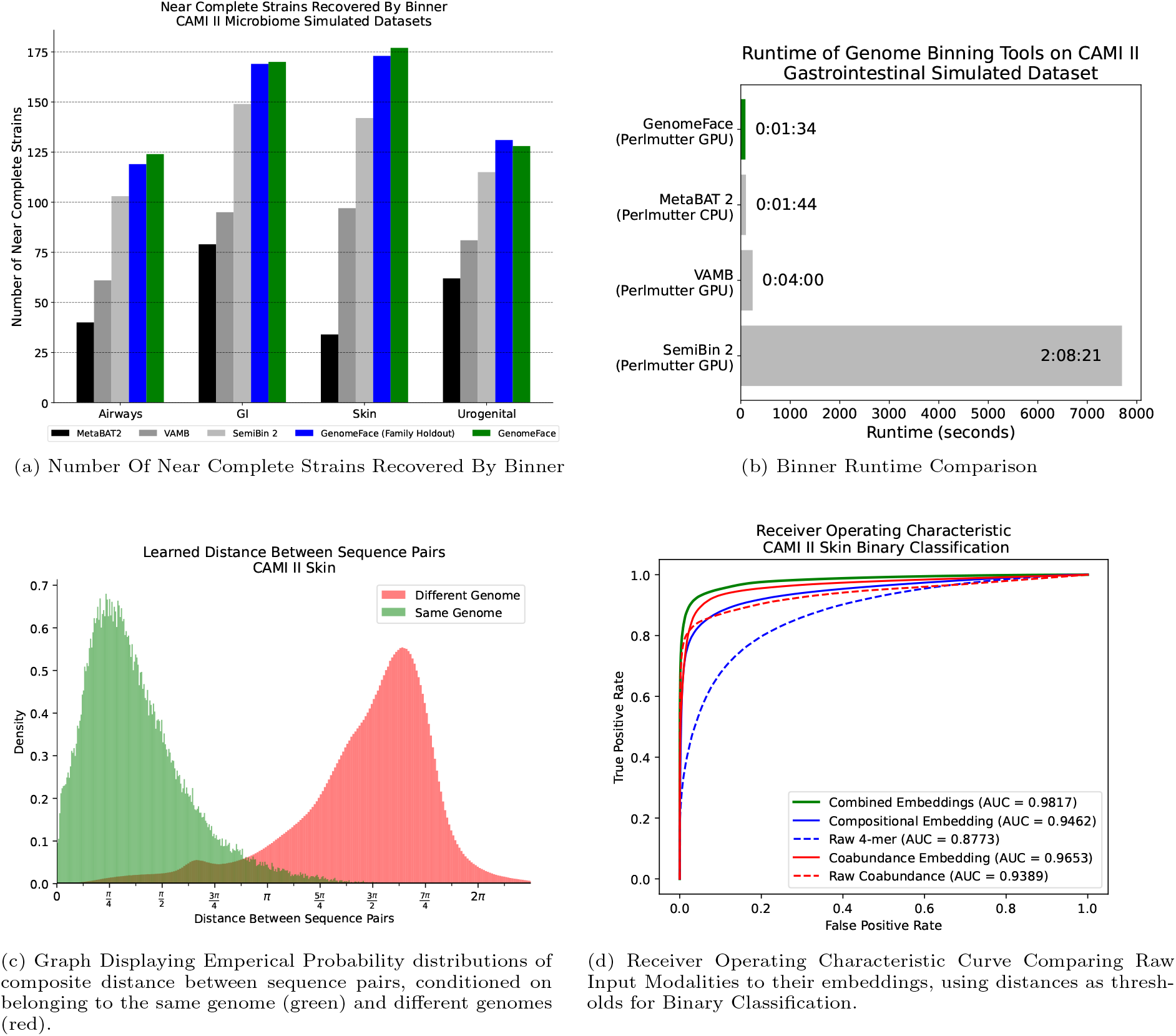
Analysis of Simulated Datasets

**Fig. 4:**
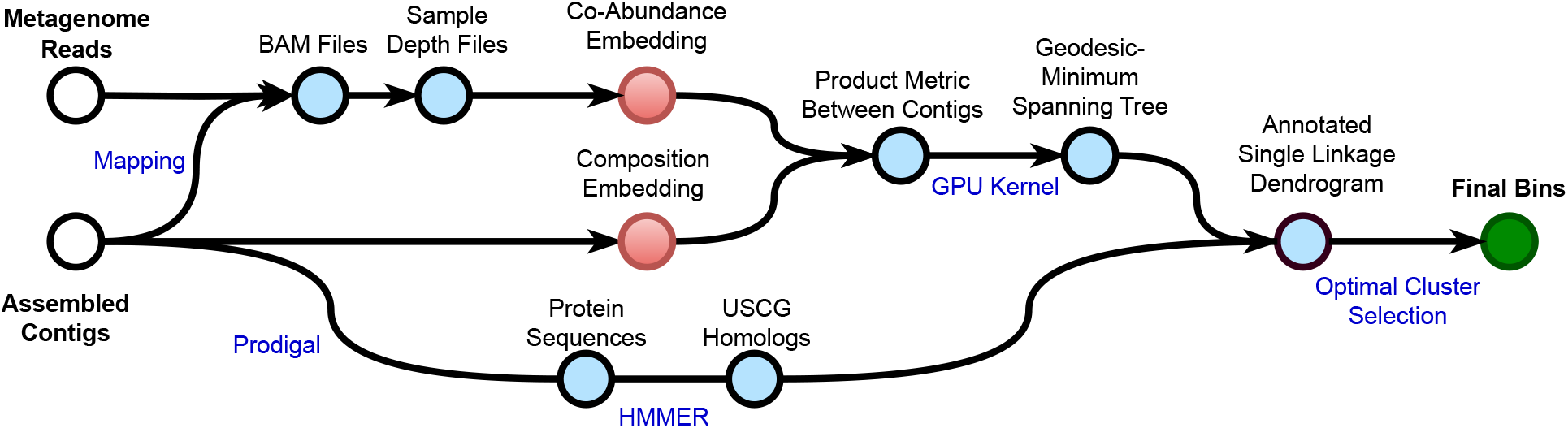
Overview of GenomeFace

**Fig. 5:**
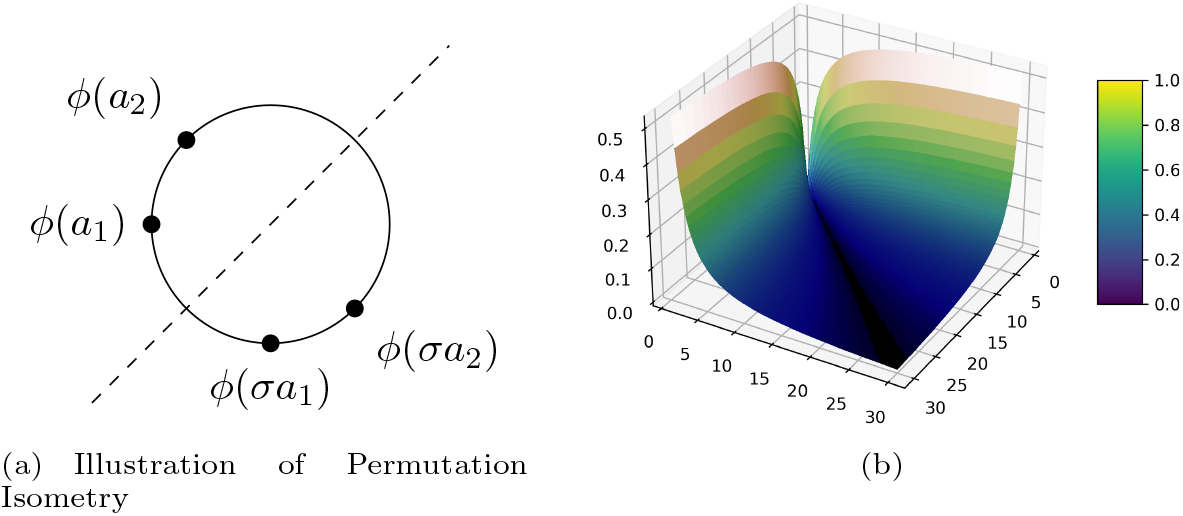
(a) Illustration of Permutation Isometry: Points *ϕ*(*a*_1_) and *ϕ*(*a*_2_) on a unit circle, and their symmetric counterparts *ϕ*(*σa*_1_) and *ϕ*(*σa*_2_) obtained through reflection across the dashed line. The isometric property of permutations is demonstrated, where distances between corresponding points remain unchanged under any permutation. (b) Surface plot illustrating the relationship between input values and the distances between embeddings in the actual trained neural network. Each point on the plot represents a pair of contigs from a theoretical single sample assembly. The parabolic shape captures the impact of read coverage values for each of the contigs (x and y axes) on the resulting distances (z axis) in the embedding space. As coverage increases, the curve gradually becomes less steep, reflecting the decreasing impact of incremental changes in coverage at higher levels, consistent with the characteristics of the Poisson distribution. This visualization demonstrates how the network approximates an implicit statistical distance, or pseudo-likelihood that contigs originate from the same genome while accounting for the underlying variability.

#### Efficiency

In all four smaller ‘toy’ datasets, GenomeFace demonstrated competitive runtimes in comparison with VAMB and MetaBAT 2. Figure 3(b) presents a comparison of the runtime of these four binners on the CAMI Gastrointestinal metagenome. The binners were benchmarked on the Perlmutter supercomputer. VAMB, Semibin 2, and GenomeFace were run on Perlmutter GPU Nodes, equipped with an AMD EPYC 7763 CPU, four NVIDIA A100 GPUs, and 256 GB of DDR4 memory. MetaBat 2, which does not utilize GPU acceleration, was allocated Perlmutter CPU nodes that feature two AMD EPYC 7763 CPUs and 512 GB of DDR4 memory, but lack GPUs.

## 3 Discussion

Microbial communities, though often dominated by a few prevalent species, are also typically characterized by a ‘long-tail’ of low-abundance organisms. Known as the rare biosphere, these organisms compose the vast majority of microbial diversity [21–24].

Despite their limited numbers, these species commonly exert disproportionate influence on their ecosystems. Often they perform specialized roles, such as autotrophic nitrifiers, or those with niche substrate preferences, such as methanotrophs and methylotrophs. Low abundance species can also play key roles, as exemplified by giant sulfur bacteria; the Beggiatoa [24, 25], Thioploca [24, 26], and Thiomargarita [24, 27] genera, while small in number, hold substantial biomass and contribute significantly to biomass and sulfur cycling in sediment due to their large-diameter cells [24, 28]. Other rare biosphere key-stone species’ presence in sediment has been shown to influence plant biomass and defensive mechanisms, having downstream effect on herbivore interactions and broader ecological dynamics [24, 29].

**Table 1:** Methodology Comparison of Binners.

| Binner | Methodology | Requires Training | Uses Marker Genes | Approx No. Params (Assume 10 Samples) |
| --- | --- | --- | --- | --- |
| MetaBAT 2 | Analytically Modeled Similarity Measure | No | No | 18 |
| VAMB | Unsupervised Embedding (Variational Autoencoder) | Yes | No | 330,000 |
| SemiBin 2 | Self-Supervised Metric Learning | Yes* | Iterative Brute-Force Reclustering | 370,000 |
| GenomeFace | Large Scale Supervised Metric Learning | No | Dynamic Programming-based Selection from Hierarchy | 100,000,000 |

Gaining insights into the rare biosphere has proven elusive, causing these scarcely understood taxa to sometimes been referred to ‘microbial dark matter’ [21]. Genome assembly in metagenomic studies has been achieved predominantly through multiassembly, a technique wherein each sample is assembled individually. While multiassembly is less computationally demanding because it processes only a subset of the data at once, this also inherently limits the effective sequencing coverage of genomes, leading to insufficient sequencing coverage for low abundance species. Multiple metagenomic studies have observed multiassembly resulting in more fragmented and incomplete Metagenome-Assembled Genomes (MAGs) displaying less diversity compared to coassembling the same data [7, 10].

This limitation is not merely anecdotal; The Critical Assessment of Metagenome Interpretation (CAMI) II challenge found in a controlled benchmark, every assembler tested required a minimum of 10 *×* sequencing coverage is necessary to achieve 90% recall of assembled genomes [18]. This implies that even with multiple rounds of multiassembly, the genomes of only the most abundant community members are likely to be constructed (and reconstructed), thereby disregarding the rare biosphere.

Coassembly offers a more comprehensive view of microbial communities, and larger coassemblies provide even greater perspective. However, this also makes the binning process more difficult; the produced assemblies are larger, more computationally challenging, and more entropically complex.

Metagenome binning, especially on datasets derived from complex environmental microbial communities, is still an unreliable process. By developing deep neural networks trained on a large corpus of high-quality reference genomes to capture sequence composition information, exploiting sample statistics to avoid sample inter-dependency, employing universal prokaryotic marker genes to guide contig clustering, and leveraging multi-GPU acceleration to enable efficient binning on extremely large datasets, we’ve shown genome binning improvements over existing software on several synthetic and real-world datasets, and in some cases, by a large margin.

### 3.1 Summary Of Contributions

#### Introduction of Massive-Scale Metric Learning to Metagenomic Analysis

Drawing inspiration from facial recognition technology in computer vision, we applied massive scale metric learning techniques to metagenomic binning, leveraging prior knowledge from tens of thousands of reference genomes in one of the most comprehensive machine learning efforts in microbial ecology. We empirically demonstrate the generalizability of these efficient techniques to distinguish between novel genomes, even at the Phyla level, advancing earlier work using.

#### Introduction of an Efficient, Structured Algorithm for Integrating Universal Single Copy Marker Genes into Clustering

Unlike other metagenome binners that use single copy marker genes in a post-clustering capacity as an oracle, GenomeFace directly incorporates these genes into the clustering process. This method constructs a hierarchy of clusters, offering a refined spectrum of sequence partitioning–from sensitive groupings to highly specific clusters. This hierarchical strategy allows for the accurate extraction of an optimal, non-redundant set of clusters in a time linear to the number of marker genes, thereby eliminating the need for multiple runs of the clustering algorithm to generate candidate clusters.

#### Binning Quality Improvements Improvements & Better Computational Performance

In addition to improving results on small-scale datasets, our techniques also enable the use of deep learning and marker genes on large-scale coassemblies with millions of sequences. This offers a significant value proposition, as large scale co-assemblies have an unparalleled ability to capture and assemble complex ecosystem’s taxonomic diversity.

## 4 Online Methods

### 4.1 Overview

To cluster assembled sequences GenomeFace uses the composition of the sequences themselves along with the per sample mapped sequencing depth. Each sequence is passed through a modality specific neural network that outputs an embedding optimized to maximize discriminative ability. Let *x*_*n*_ represent the output of the composition based embedding for the nth sequence, and *y*_*n*_ for co-abundance based embedding. The distance between two sequences **C**_*i*_, **C**_*j*_ is then defined based on their embeddings as

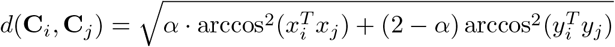

where α ∈ [0, 1] is a mixing coeffient to weight the contribution of each modality, that (by default) is determined by the number of environmental samples.

We consider each sequence a vertex in a graph, and find the minimum spanning tree of the complete graph where the edge weight between c_*i*_ and c_*j*_ is defined by the distance between them using our GPU accelerated kernel. We term this graph the ‘geodesic minimum spanning tree’ or G-MST.

From the G-MST of N sequences, a single linkage dendrogram based on the distances is built to form a hierarchy of 2N *−* 1 *candidate* clusters. Each leaf cluster **C**_*n*_ xx‘is annotated using the Universal Single Copy Genes the sequence it represents codes. These genes are propagated upward in the dendrogram to obtain marker gene counts for every candidate cluster based on the sequences they contain, and bosed on their annotated genes. An abstract reward value is then calculated for each candidate cluster based on their gene counts, and an arrangement of candidate clusters, **Clusters**, is outputted to maximize 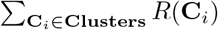 based on a cluster reward function *R*.

### 4.2 Compositional Neural Network

#### 4.2.1 Pipelined Contig Sampling and Feature Generation

To train our compositional neural network genomes from the Genome Taxonomy Database were filtered to those with an estimated CheckM contamination of less 5%, and a minimum completion of 80%. The minimum completion requirement was implemented out of concern the network would incorrectly bias homologous regions of a genomes to a certain taxonomy. A limit of 25 genomes per genera was implemented, as to satisfy memory requirements and prevent severe taxonomic imbalances in our training set. This results in 43,096 genomes for our primary training.

A native python module was developed as to be called by a Tensorflow Dataset Iterator. The python loads a collection of genomes into memory, and calls a sample method. When the sample method is called, a batch of 2^18^ genomes are chosen, with replacement, as to generate a training contig from. For each sampled reference genome, a scaffold is chose from those with probability proportional to the length of the scaffold. On this scaffold, a window of length 2000 is sampled uniformly to represent a contig. For these 2^18^ contigs, their 1-5-mers frequencies are counted, with reverse complements considered equivalent, such that for *k*-merse of even *k*, a feature vector of length 2^*k−*1^ + 2^2*k−*1^ is formed, and of length 2^2*k−*1^ for odd *k*-mers. For tractability, the degenerate R-Y alphabet where A,G ↦ R, and C,T ↦ Y, *k*-mers of length 6-10 are used with reverse compliments being considered equivalent, such that for even and odd length *k*, feature vectors of size 2^*k*/2*−*1^ + 2^*k−*1^ and 2^*k−*1^ were respectively produced. After counting, each respective kmer frequency vector was *l*_1_ normalized.

We parallelize *k*-mer counting of each batch with multithreading and during the processing the Python Global Interpret Lock (GIL) is released. This is safe, as no Python managed memory allocations are accessed or made. After frequency counting is complete, the GIL is reacquired, the kmer frequencies are coerced into a tensorflow friendly input by using the NumPy C-API, and returned to Python with labels.

By instructing Tensorflow is instructed to prefetch batches, we are able to asynchronously sample and count kmers for sequences in for one batch utilizing the CPU, while simultaneously training the network using the GPUs in our system.

#### 4.2.2 Network Architecture

The input to the composition neural network is the 1-5mer and 6-10RY-mers feature vectors, as described above. These are concatenated to form one logical vector feature vector. Since we sample contigs, it is non trivial to normalize the ‘dataset’ in its entirety, so a Batch Normalization layer is applied to this concatenation. A dense layer with output size 4096 with hyperbolic tangent activation is applied, then another Batch Normalization layer. Again, another a dense layer with output size 16384 and a hyperbolic tangent activation follows, and another Batch Normalization Layer follows that.

Similarly to described in [insert paper here], we form our embedding by applying a Dense layer with output size 512. No (i.e. a “linear”) activation is applied, followed by a batch normalization layer. The output of this batch normalization layer is then *l*_2_ normalized, completing the base embedding network. For regularization, weights of every dense layer has a very mild *l*_2_ penalty applied.

### 4.3 Co-Abundance Network

Like other metagenome binners, we integrate per sample read coverage into our binner. Disregarding sequencing bias, reads in single environmental sample from a genome can be seen as being drawn at uniform rate throughout the genome – thus the mean read coverage for a contig can be seen as a proxy measure for abundance of the organism from which it originates, with each sample providing additional information for distinguishing between genomes in the binning process. Viewed as a vector, the mean per sample read coverage could of course be clustered on directly. However, this is less than ideal. The uniform coverage rate assumption implies the distribution of coverage for per base pair in a genome is theoretically Poisson, Thus as read coverage increases for a genome, we would expect mean variance does also, indicating a constant differential in read coverage between two contigs should be viewed less significantly at a higher coverage than a lower one. In observation, the variance of the coverage is significantly greater what is expected of the Poisson distribution, due to factors such as sequencing biases and read mapping ambiguities.

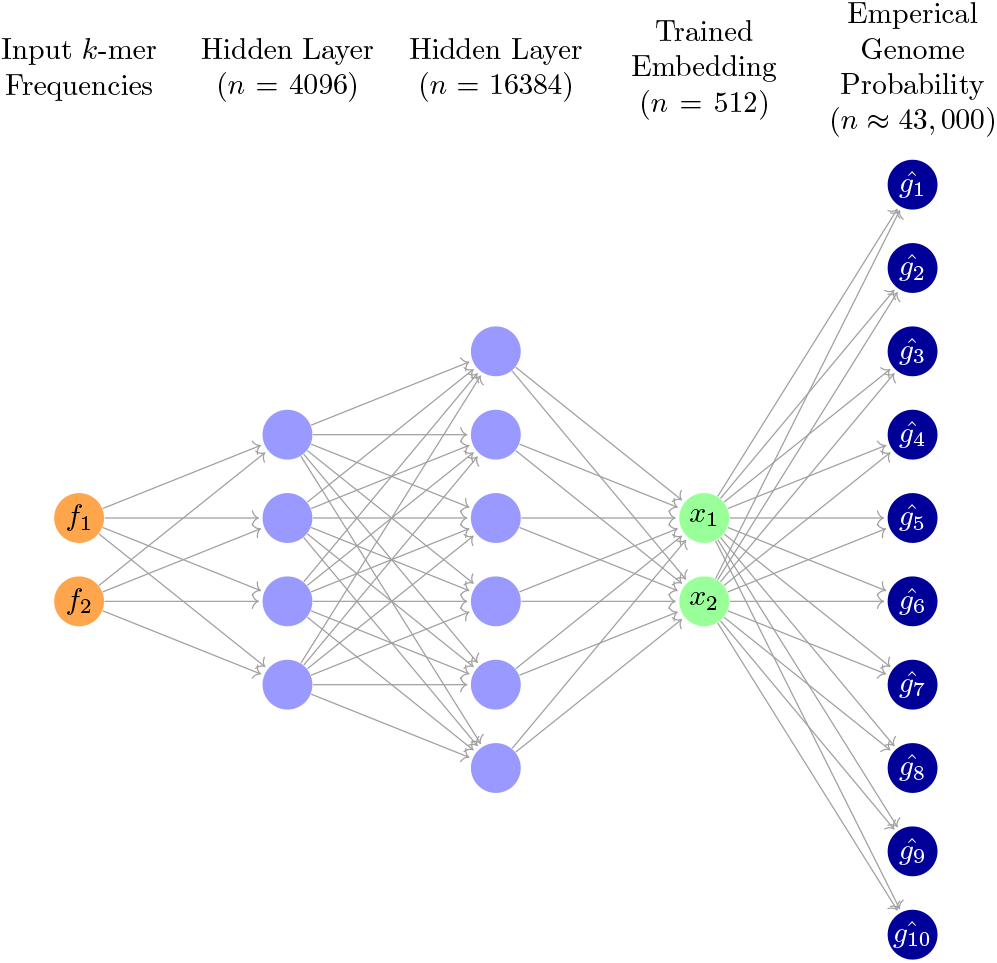

To account for this, we train a second network to embed our coverages into a higher dimensional space. However, as the number of environmental samples used for a ecological study can vary from as few as 1 to more than 1000, the appropriate construction of a general network to do so is not immediately obvious. Additionally, since each sample depth coverage portrays the same type of information, there is some symmetry inherent to the problem.

Making clear this symmetry, we design our Co-abundance neural network to enforce the principle of permutation invariance in our output binning: the order of input of the environmental sample coverage does not affect the produced clustering. Before we describe the implementation, we will justify its properties.

The motivation for permutation invariance arises from the view of the samples as exchangeable from a Bayesian perspective. In statistics, a sequence of random variables (in our case, the samples) is considered exchangeable if its probability distribution remains unchanged when the variables are permuted. Although each sample may come from a unique source or biome, from the perspective of the binner, the order does not convey any semantical information that should influence the output. Therefore, for robustness, these samples should be treated as exchangeable, operating under the premise that any given order of samples could have occurred under the underlying probability distribution.

We denote our abundance embedding network by *ϕ* : ℝ^|*samples*|^ → ℝ^*m*^. As clustering algorithms operate on distances, the idea of permutation invariance in the produced binning translates to an intriguing property for our neural network: for any two abundance vectors a_1_, a_2_, the distances in the embedding space should remain unchanged under any permutation of the samples. Formally, for every permutation *σ*_*i*_, we have

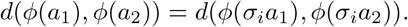

This observation leads us to a key conclusion: for every permutation *σ*_*i*_, we can find a corresponding transformation *U*_*i*_ : ℝ^*m*^ → ℝ^*m*^ such that *U*_*i*_ ◦ *ϕ*(*a*) = *ϕ*(*σ*_*i*_*a*). The transformations *U*_*i*_ form a group that is isomorphic to the group of permutations. Specifically, the group structure is induced by each *U*_*i*_ being an isometry, i.e., it preserves distances in ℝ^*m*^. This characterizes our neural network *ϕ* as an equivariant map, and that it should “pass” the permutation group through it, and enforcing a factorial way symmetry (i.e. |sample|!) on it’s input. To enforce this on the symmetry, we use a permutationally equivariant neural network.

One might naturally question the benefit of a permutationally equivariant neural network, as opposed to just mapping each sample to a higher-dimensional space independently and then concatenating the results. Crucially, our approach recognizes that the sequencing depths across different samples are not independent - they exhibit systematic biases due to sequencing techniques and factors like sequence composition and position within the genome. Moreover, our model allows for a representation that avoids the naive averaging, instead allowing for a geometry which could more accurately describes the joint likelihood of the differential abundances across multiple environmental samples.

#### 4.3.1 Co-Abundance Network Architecture

To construct the coabundance network, we employ Set Attention Blocks from Set Transformers, with modifications including the removal of layer normalization and alteration of the activation function in the Feed-Forward Network (FF) to the hyperbolic tangent function. For ease of explanation, the batch dimension is omitted.

The process begins by generating the initial matrix *M*_1_ through the outer product of d, the per sample mean sequencing depth of a contig, with a learnable weight vector v. This matrix, along with the subsequent matrices *M*_2_ to *M*_5_, are of size NumSamples *×* 16 and have a rank of 2. These matrices are iteratively processed through Permutation Equivariant Blocks, each of which consists of a multihead self-attention mechanism followed by a Feed Forward (FF) network. These block enhances the feature representation of each sample while maintaining the permutation equivariance peroperty in the first index of the matrix, which corresponds to the individual sequencing sample.

After the series of permutation-equivariant transformations, the matrix *M*_5_ is flattened to form *M*_6_, changing its dimensions to NumSamples *·* 16 and reducing its rank to 1. The flattened matrix is then *l*_2_ normalized (in *M*_7_) on to the hypersphere.

##### Algorithm 1

Co-Abundance Network

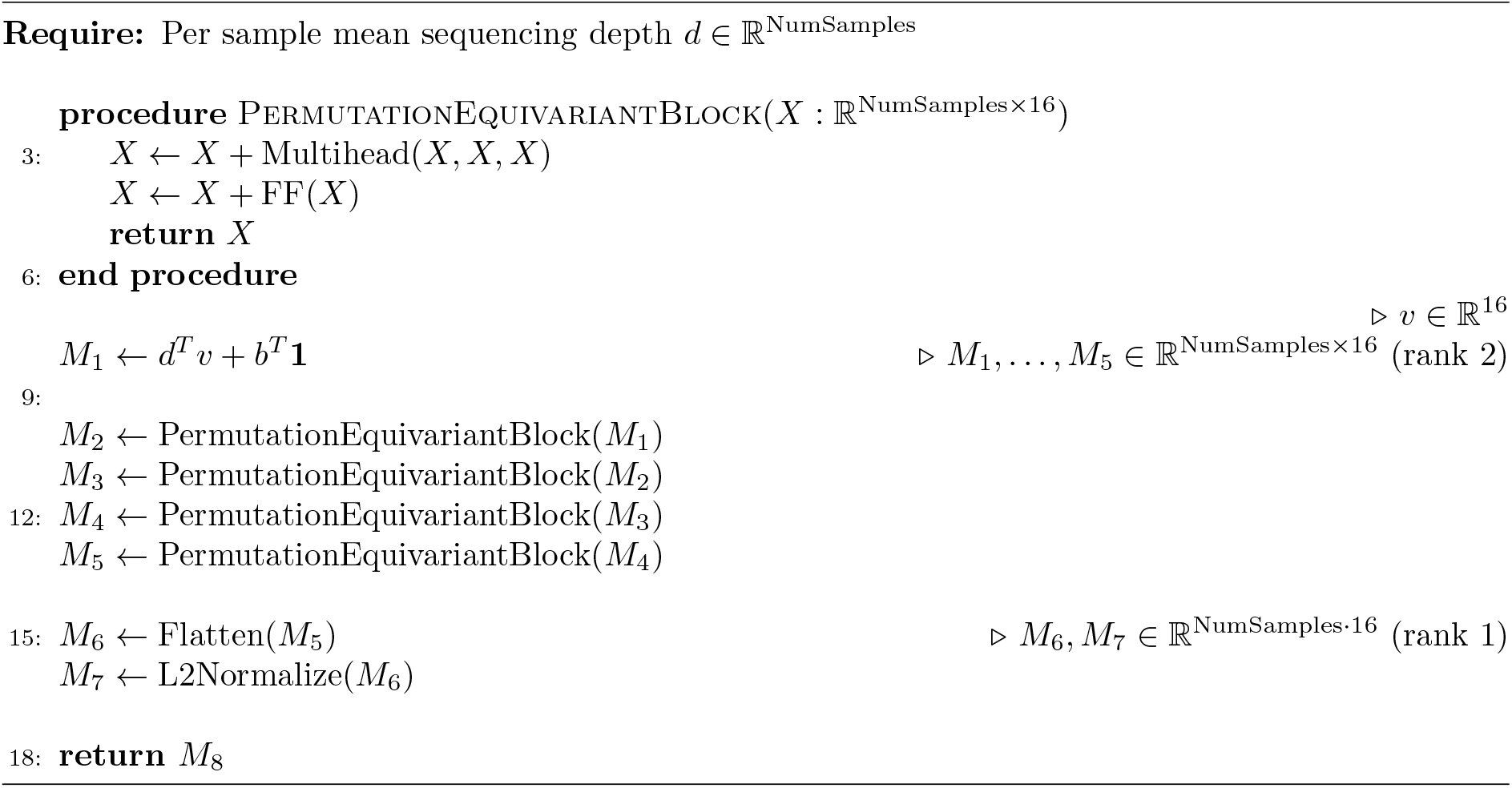

#### 4.3.2 Training the Networks

The popular softmax loss often comes into play in deep neural networks when addressing multi-class classification challenges. It is the combination of the softmax activation function in the network’s output layer and then proceeds with categorical cross-entropy training. In the following passage, we detail the intuition for is use in our embedding training.

In a network engineered for *d*-way categorical classification that employs softmax loss, the second to last layer produces an embedding of size *n*. The final layer then multiplies this embedding by a matrix W, which exists in ℝ^*d×n*^, consequently yielding a *d*-dimensional vector. Each vector component aligns with a class. The softmax activation guarantees that the vector components resulting are not only positive but also sum up to one, thus producing a valid predicted probability distribution.

We’ll refer to the *a*-th row vector of the matrix *W* as *W*_*a*_. Our network employs the following equation to estimate the probability of a contig pertaining to genome *k*:

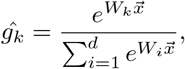

where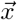 is the sample’s embedding from the second-to-last layer of the network.

The network is trained by minimizing the cross-entropy *H* between the predicted and ground truth probability distributions. Cross-entropy quantifies the minimum average number of bits required to represent events from distribution *p* using an optimal coding scheme designed for distribution *q*. It is defined as:

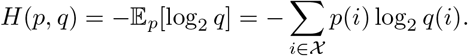

Assume *y* represents the ground truth distribution, embodied as a one-hot vector, with *k* indicating the true class of the sample. We can express the softmax loss as follows:

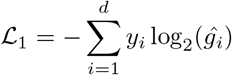

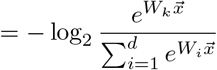

Previous studies noted that we can interpret the output from the second to last layer as distributing classes radially in a natural manner. Observe that for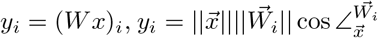. In this context, the magnitude of the output vector may suggest the confidence level of the classification. However, it’s the radial angle between the output vector and the weight vectors, in conjunction with their relative magnitudes, that naturally encodes the ‘feature description’.

To further enhance the discriminative capacity of the features, we apply *l*_2_ normalization to both the rows of *W* and each 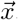. This normalization ensures that their product, and hence the per-class probability, depends exclusively on the angle between them. Following this, we introduce a softmax scale (or inverse-temperature) denoted by *s*.

Under this construction, each component (*W x*)_*i*_ of the matrix-vector product 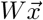 can be interpreted as the cosine of the angle between the *i*th row of *W* and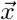. From a heuristic viewpoint, we can consider the *i*th row of W as the ”center” embedding that represents its corresponding class. With these modifications, the loss function can be reformulated as:

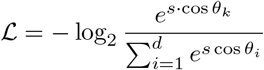

Minimizing the cross-entropy loss implies that, for each sample, the embedding is close to its class center (θ_*k*_ is small) and far away from the centers of other classes (*θ*_*i*_, *i* ≠ *k* is large). To further enhance the model, a softmax scale (or inverse-temperature) s is applied. This increases the range of logits entering the softmax function, specifically the cosines. For large *d*, without the scale factor *s*, the max achievable probability would be limited to at most *e*/*d* in the ideal case.

While this captures the essence of our network, we integrate the modificaitons to softmax-crossentropy presented by CurricularFace [30] for curricular learning. This revised loss function is:

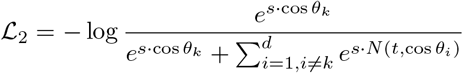

Here, *N* is defined b:

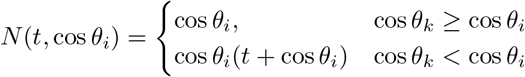

Where *t* represents the exponentially weighted average of cos *θ*_*k*_ across batches.

### 4.4 Balancing Embeddings

Let *x*_*k*_, *y*_*k*_ respectively denote the embeddings produced by our compositional and coabundance networks for a sequence *c*_*k*_. We define the composite distance between two sequences *c*_*i*_, *c*_*j*_ at the

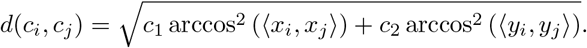

where *c*_*1*_, *c*_*2*_ are mixing coeffient determined by the number of samples.

#### Default Method

We model the weight placed on each modality should be inversely proportionally to the inter-class variance of the modality. We assuming the variance of the compositional embedding, and therefore *c*_*1*_, should be constant with regards to number of environmental samples *n*.

For the Coabundance embedding, we modeled the a variance component (*𝓋*_*1*_) term which should be inversely proportional to the number of samples, as though they were independent, however, considering systemic biases (such as read mapping ambiguities caused by repeated regions) which would be constant (*𝓋*_*2*_). Therefore, we arrived that *c*_*2*_, reciprocal to the variance, would be

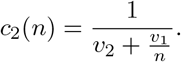

To determine the constants *𝓋*_*1*_, *𝓋*_*2*_ we took the follow steps:

1. We simplified the dissimilarity formula by setting *c*_1_ to a constant value of 1. This adjustment is based on the understanding that scaling both *c*_1_ and *c*_2_ by the same factor only scales the output scores by a constant, leaving the AUROC unchanged.
2. We considered the effect of using various number of environmental samples *n* on the output embedding with using the CAMI II Airways dataset by only considering the first *n* samples, therafter removing sequences with no coverage after modification. By inspection, it was determined the AUROC of the composite distance as a function of c_2_ was a unimodal function. Thus, for each number of samples *n*, we used golden section search to find the value of *c*_2_ that maximized the AUROC for the restricted dataset. This resulted in a series of pairs (*n, c*_2_(*n*)) correlating each sample size with its corresponding optimal *c*_2_ value.
3. Finally, we used the pairs of (*n, c*_2_(*n*)) as data points for curve fitting. Applying Scikit-Learn’s curve fit tool, we fit the function *c*_2_(*n*) defined above, allowing us to fit *𝓋*_1_, *𝓋*_2_ to the observed pairs, thereby allowing us to predict the optimal *c*_2_(*n*) for any number of samples *n*.

After, to standardize the range of our dissimilarity metric to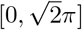 we scale *c*_1_ and *c*_2_ such that *c*_1_ + *c*_2_ = 2.

#### Mixing using Bayesian Optimization

We present the option to balance the two embeddings automatically adaptively, by repeatably clustering on a subset of the sequences. We define *c*_2_ = 2 *− c*_1_, then fit c_1_ using Bayesian Optimization to optimize the total cluster reward, as defined in the next section, by repeatedly clustering on the largest 200,000 sequences. After an optimal *c*_1_ value is found, all of the sequences are clustered.

### 4.5 Clustering

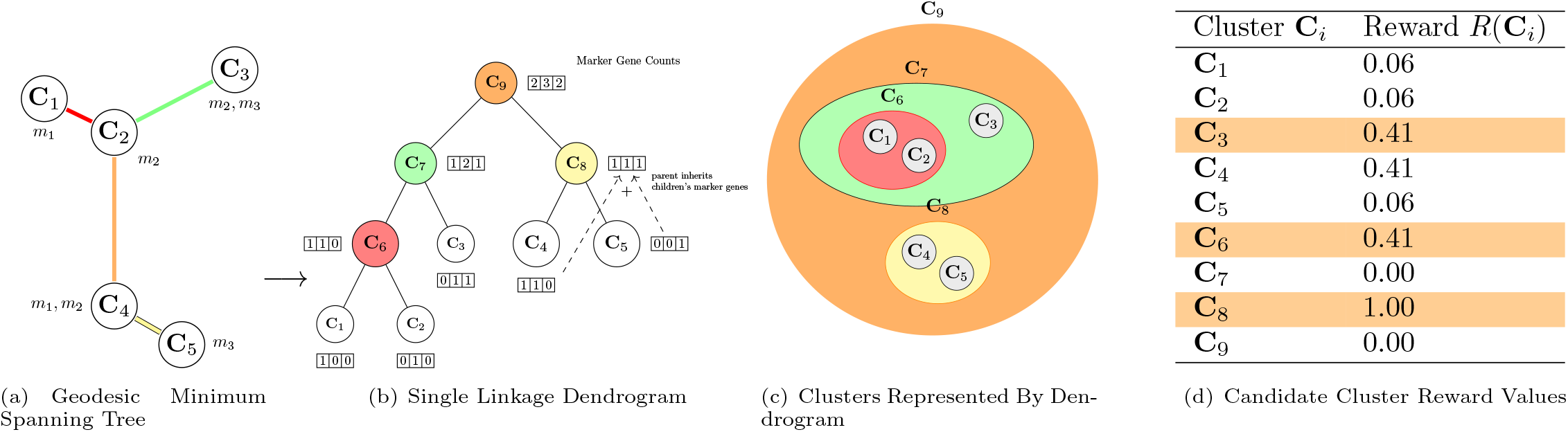

Our clustering algorithm leverages domain-specific knowledge to achieve more accurate and meaningful cluster assignments. Universal Single Copy Marker Genes (USCG) can be viewed as weak pairwise labels. Two occurrences of the same USCG imply that the contigs coding for them should be in different clusters, while we know that each cluster should ideally include one of each gene. By incorporating information from universal single-copy marker genes, our method can make more informed decisions when merging or separating clusters compared to generic clustering approaches that rely purely on distance or density estimates alone. Our tailored approach to clustering provides a distinct advantage over traditional, non-task-specific clustering algorithms and ensemble binners, allowing for improved overall quality of the resulting genomic bins.

The algorithm operates by first building a Geodesic Minimum Spanning Tree (G-MST) from the contigs’ embeddings. It then constructs a Single Linkage Cluster Dendrogram from the G-MST, which serves as a hierarchical representation of candidate clusters. The cluster represented by each parent node is the merger of the clusters represented by its child nodes, where the leaf nodes represent clusters defined by a single contig. This hierarchical representation allows us to efficiently incorporate marker gene information without the need to reprocess contigs or marker genes for every candidate cluster considered. By simply adding the marker gene counts of child nodes in the dendrogram to obtain the marker gene counts of each parent node, we can quickly estimate the purity and completeness of each candidate.

Additionally, this hierarchical structure presents a topological ordering on the candidate clusters. This topological ordering enables us to find the optimal set of clusters from the candidate clusters in near linear time by using dynamic programming. In contrast, ensemble binners face a far more challenging task—an NP-complete problem—as finding the optimal set of clusters for an arbitrary set of candidate clusters essentially involves solving an optimal weighted set packing problem, where the weight for each set is the cluster’s reward value.

Our algorithm can be seen as examining balls in the embedding space surrounding each contig’s embeddings, which form a cover for the contigs’ embeddings. Contigs clusters are naturally defined by the connected components of this cover. Nodes in the dendrogram represent thresholds such that the connected components have changed. As we navigate the dendrogram, the balls either expand or contract based on the edge weights of the dendrogram, as induced by the Geodesic Minimum Spanning Tree.

By integrating marker gene information directly into the clustering process, our algorithm learns in a transductive or semi-supervised manner from the data. The marker genes serve as weak pairwise labels, supplying information about the relationships between contigs’ classes. This information guides the decision-making process for growing or shrinking the balls in the embedding space. If the presence of marker genes signals potential contamination in a cluster, the algorithm may opt to shrink the balls around the contigs that cluster as to fracture it and lower contamination; on the other hand, if the marker gene information suggests that merging clusters would improve completeness, the algorithm may choose to grow the balls surrounding the pertinent contigs. We actively learn a non parametric decision boundary for inclusion for each cluster – not only utilizing those contigs with marker genes, but also those without. For further intuition, refer to HDBSCAN’s [31] paper, from which our algorithm was based.

Formally, given the clusters represented by taking the nodes of the Single Linkage Dendrogram, and R defined to be a function that outputs a real valued rewards a cluster based on it’s marker gene contents, we optimize to find a set of clusters from the hierarchy **Cluster**

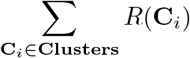

subject to the constraint that **Clusters** is a partition of the contigs – every contig exists in some cluster, and no contig is in two clusters.

We give an overview of the algorithm, detail our chosen cluster reward function, and then give the algorithm in detail.

#### 4.5.1 Overview of the Clustering Algorithm

The main steps of the clustering algorithm are as follows:

1. Build the Geodesic Minimum Spanning Tree (G-MST) from the contigs’ embeddings.
2. Construct a Single Linkage Cluster Dendrogram from the G-MST and initialize a marker gene count for each node in the dendrogram.
3. Enumerate and process marker genes for each contig.
4. Traverse the dendrogram in reverse topological order, propagating marker genes counts upwards, and computing the reward for each node.
5. Traverse the dendrogram in topological order to extract the final set of clusters, choosing the optimal reward clusters.

#### 4.5.2 Cluster Reward Function

Following the premise that universal single-copy genes (USCGs) should occur in every genome and be single-copy, we use the frequency of their occurrence (or over-occurrence) to measure the completion and contamination of a cluster.

In our tests, we used the set of 40 USCGs from Creevey et al [32]. that were derived from Bacteria, Archaea, and Eukaryotes for their generality and were evaluated to be of higher quality in terms of measured universality and uniqueness compared to other gene sets[33]. Similarly to CheckM1’s methodology of grading the first marker gene as correct and viewing any later occurrences as contamination[34], we define the following heuristics to estimate Purity and *F*_1_ score. Let *M* be our set of universal single-copy marker genes, and for *m ∈ M*, define Count_*m*_(**C**_*i*_) to be the number of times the marker gene m occurs in some cluster **C**_*i*_. Then

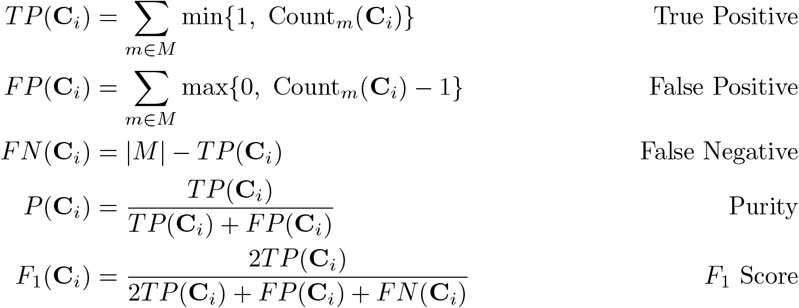

While any cluster reward function may be chosen, we choose the reward function

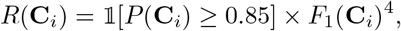

as a good trade-off between valuing purity and completeness. We use the *F*_1_ score rather than the *F*_*β*_ with a lower *β* to be more tolerant of genomes with mutations where a small number of genes in our set become non-single copy, but compensate with a default minimum 85% estimated purity for any reward (modifiable as a tuning parameter). Since the raw *F*_1_ score is subadditive with respect to the merger of two bins, even if there was no apparent contamination, this would result in multiple small incomplete bins being preferred to a larger completely pure one. To negate this, we map the *F*_1_ score through a sufficiently convex function, and arbitrarily choose to raise it to the fourth power.

##### Geodesic Minimum Spanning Tree Construction

We modify NVIDIA’s [35] HDBSCAN implementation for euclidean minimum spanning tree construction which finds the *k*-nearest nearbors using FAISS’s *k*-select [36] fused with a matrix multiply kernel to calculate the euclidean *k*-nearest neighbors, thereafter finding the Euclidean-Minimum Spanning Tree [37].

NVIDIA’s implementation to find the euclidean distance by combining with magnitude of input vectors with their respective inner products. It can be conceptualized as loading the vectors, denoted *𝓋*_*n*_ as the rows matrix V, and finding *V V* ^*T*^, then calculating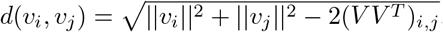. (This is done interspersed with reductions with the *k*-select, the actual calculation of (*V V* ^*T*^) in it’s entirety would be prohibitively large.)

We instead use two matrices, denote *X* and *Y*, one for each modality, such that the *n*th row of each represents the embedding of the *n*th sequence for that modality. The two matrix multiplies are performed sequentially the first pipelined into the second, then the k-select is applied in a similar manner as previously, using the two matrix multiplies to calculate the distance as *d*(*C*_*i*_, *C*_*j*_) = α arccos^2^((*XX*^*T*^)_*i,j*_)+ (1 *−* α) arccos^2^((*YY* ^*T*^)_*i,j*_) for some mixing cosntant α.

The kernel was also modified to utilize all available GPUs when finding the *k*-nearest neighbors.

##### Efficient Single Linkage Dendrogram Construction

To ensure clarity, we detail the process of constructing the Single Linkage Dendrogram using the weight-sorted edges of the Geodesic Minimum Spanning Tree (G-MST). This construction can be efficiently achieved in Nα(N) time using a modified union-find data structure, where N represents the number of sequences to be clustered, and α is the inverse Ackermann function, which is effectively constant for all practical purposes.

We introduce a modified version of the union-find data structure. This enhanced structure allows each set to reference a vertex in our dendrogram. For a given set s, this reference field is denoted as s.ref.

The initial traversal of the dendrogram following its construction can be omitted. Since the dendrogram is built in reverse topological order relative to the root, the steps usually performed during the first traversal can be integrated into the construction phase itself. However, for simplicity, we describe these as separate steps in our presentation.

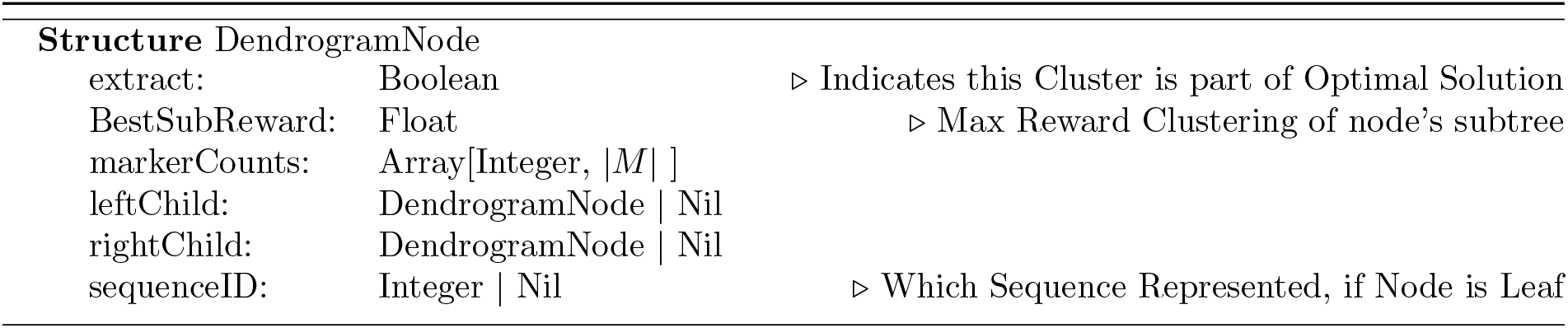

###### Algorithm 2

Efficient Single Linkage Dendrogram Construction.

Each leaf node in the dendrogram is initialized with a ‘markerCounts’ vector, where each index corresponds to a specific gene, and the value at that index indicates the count of occurrences of the gene in
the associated sequence.

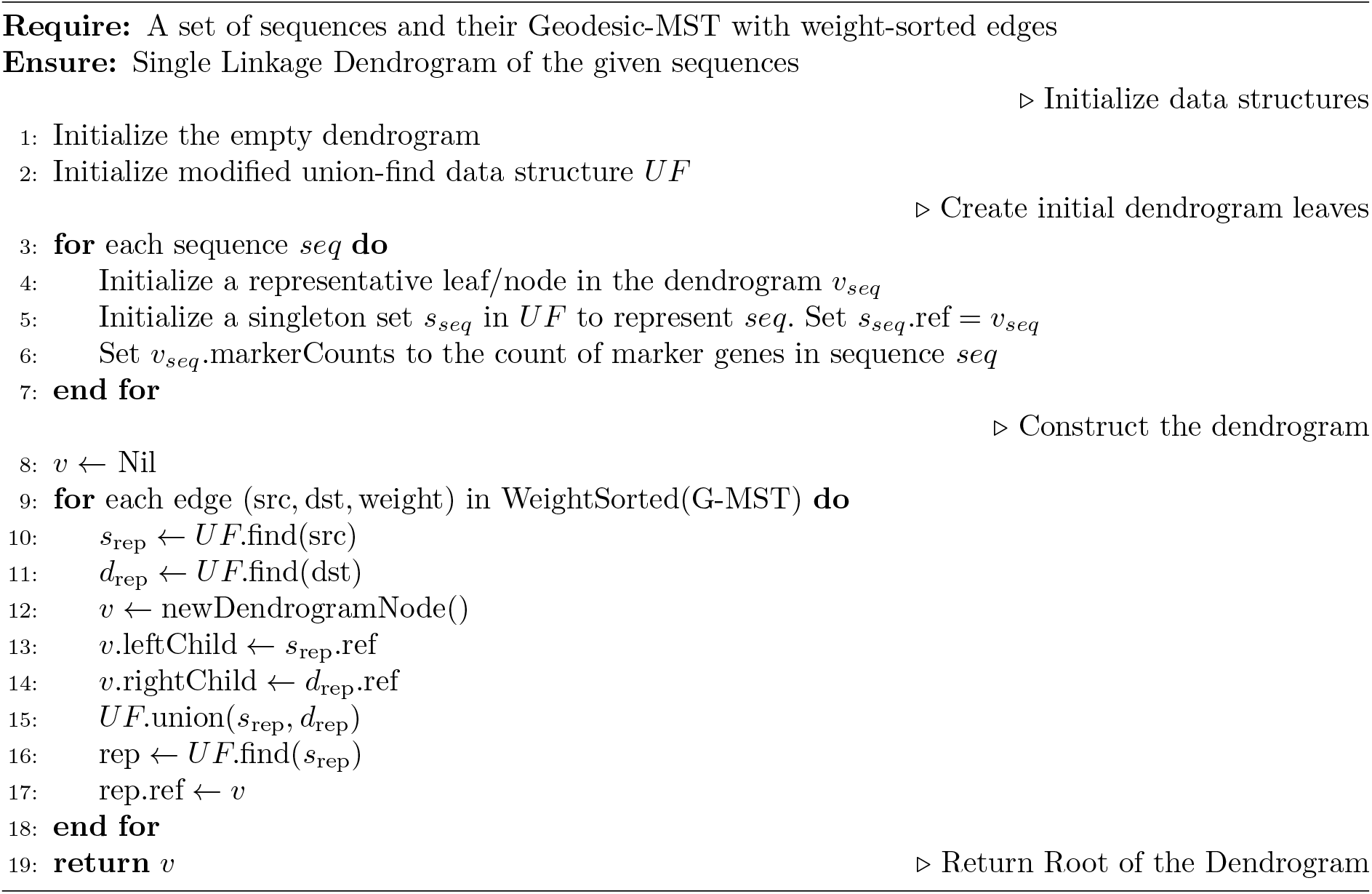

##### Marker Gene Propagation and Cluster Optimization

In this step, we use dynamic programming to identify the optimal set of clusters, **BinOutput**, that given cluster reward function R maximizes the sum of rewards,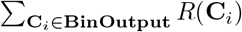, under the constraint that **BinOutput** forms a partition of the input sequences, i.e. each sequence belongs in exactly one output cluster. Here, **C**_*i*_ represents nodes in the dendrogram, with clusters defined by the points representing their leaves.

This is achieved by solving the subproblem defined for a node **C**_*i*_, the maximum reward achievable considering only the clusters within its subtree for inclusion in **BinOutput**. The base case for leaf nodes is direct, where the reward is *R*(**C**_*i*_). For non-leaf nodes, we compute the maximum reward by comparing the reward for including the node itself in **BinOutput**, *R*(**C**_*i*_), against the sum of the optimal rewards from its child nodes’ subtrees. This is because by our partition constraint, if some node **C**_*i*_ ∈ **BinOutput**, none of its descendants in the dendrogram can be, as they represent members of refinements of **C**_*i*_. Thus, traversal is performed in reverse topological order, allowing us to calculate each node’s **C**_*i*_.BestSubReward up to the root, which yields the maximum reward for the entire sequence set.

The optimal set **BinOutput** is then extracted in the next step by a forward traversal, selecting clusters marked for extraction using the extract field on the dendrogram nodes, and collecting the sequences their leaves represent.

Note, when we set **C**_*i*_.extract as True, we do not remove the flag from their descendants. Doing so would increase the complexity. This is of no concern, as we may simply stop traversal of **C**_*i*_’s descendants during our next step.

###### Algorithm 3

Marker Gene Propagation and Reward Calculation

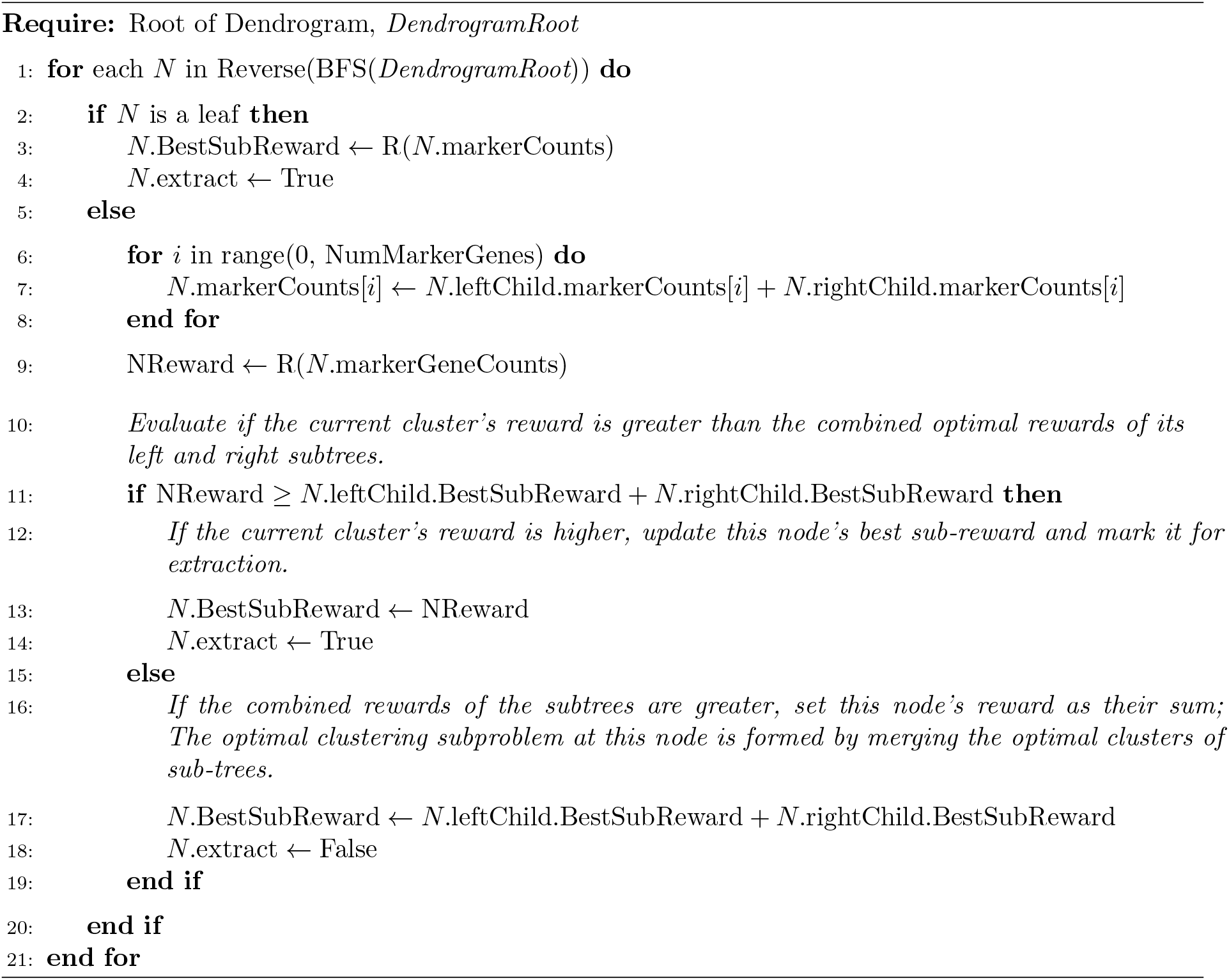

##### Cluster Extraction

For completeness, we detail how to extract the clusters from the dendrogram after Cluster optimization is performed in algorithm 4.

###### Algorithm 4

Cluster Extraction

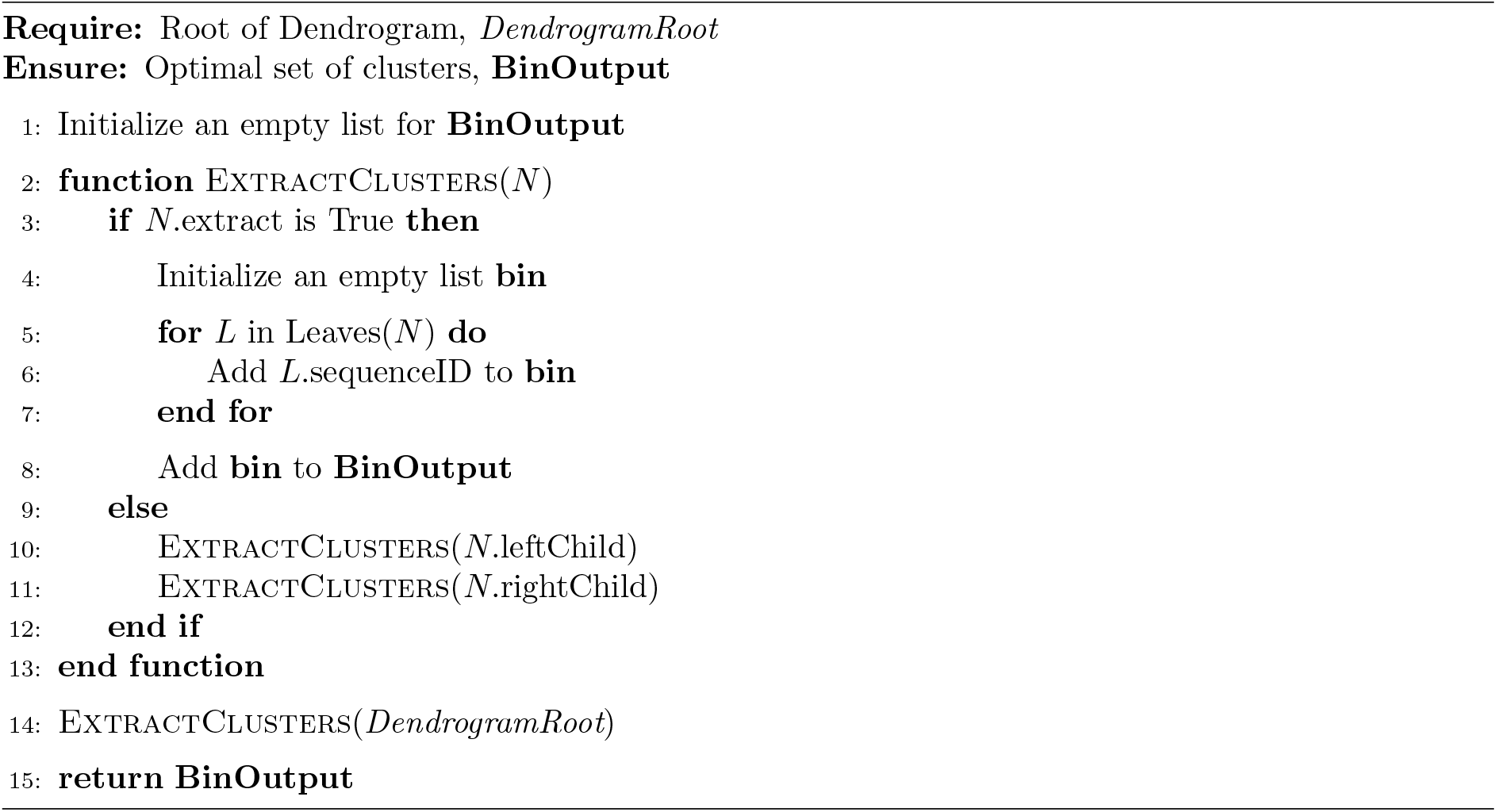

### 4.6 Additional Sythethic Dataset Evaluation Details

For the evaluation of the CAMI Human Microbiome datasets, we recreated similar benchmarks to those in VAMB and Semibin 2’s papers. The reference, or ’gold-standard’, assemblies and simulated reads for each sample were acquired from the CAMI challenge resources [**?** ]. We combined the gold-standard assemblies from each collection site into a single fasta ‘multi-assembly’ file. Subsequently, the simulated read sets corresponding to these assemblies were aligned using BWA-MEM [38]. This alignment process yielded BAM files. Coverage profiles in a format compatible with MetaBAT 2 were generated using CoverM, and these profiles were used as input for MetaBAT 2, VAMB, and GenomeFace. Semibin 2 was provided with the raw BAM files directly, as it is incompatible with the MetaBAT 2 coverage profile format. We used VAMB’s benchmarking tool in our evaluation, which measures the purity and completeness of each genome bin in relation to the number of base pairs in the gold-standard assembly. This tool adopts the same evaluation methodology as the AMBER benchmarker [20], which was used in the CAMI challenge. Since the multiassembly produced from Oral site samples were used used to train the co-abundance network, it was excluded from evaluation.

## Declarations

- The work conducted by the U.S. Department of Energy Joint Genome Institute (https://ror.org/04xm1d337), a DOE Office of Science User Facility, is supported by the Office of Science of the U.S. Department of Energy operated under Contract No. DE-AC02-05CH11231.
- Authors’ contributions
  – Richard Lettich and Katherine Yelick conceived the study.
  – Richard Lettich designed the clustering algorithim, trained the neural network, modified the GPU clustering Kernel and wrote the GenomeFace program.
  – Richard Lettich, Robert Riley, Zhong Wang wrote the paper
  – Andrew Tritt, Robert Egan, and Richard Lettich Collected training data.
  – Richard Lettich performed simulated dataset prepossessing.
  – Robert Egan and Robert Riley performed Real Dataset preprocessing and Assembly.
  – All authors participated in planning and analysis of real dataset Case studies, reviewed the manuscript, and approved the final version (pending).

